# Classification with Missing Data - A *NIFty* Pipeline for Single-Cell Proteomics

**DOI:** 10.64898/2026.03.06.710179

**Authors:** Alyssa A Nitz, Blake McGee, Benjamin Echarry, Samuel H Payne

## Abstract

Single-cell proteomics (SCP) is uniquely suited for cell-type characterization, trajectory-based inference, and microenvironment mapping. Evaluating biological hypotheses in these experiments requires labeled cells. Without a pre-measurement label, machine learning is used to identify features that characterize the cell types and classify unlabeled samples. Current implementations of annotation methods come with several statistical and computational disadvantages. First, machine-learning methods require complete data, leading to large amounts of missing-value imputation in SCP. Additionally, some machine-learning methods select features and classify samples via cross-sample comparisons, nullifying downstream cross-sample comparisons, like differential expression, through double dipping. Finally, measurements from different proteomic experiments are not directly comparable due to batch effects, significantly limiting the accuracy of classifiers trained on external data. Here we present NIFty, a top-scoring pairs based feature selection method, implemented in a full classification pipeline, that does not require pre-imputed data as input or employ circular analysis techniques, and overcomes batch effects without batch correction. When tested on imputed vs unimputed data, data with large batch effects, and multiclass data, NIFty successfully overcame the targeted classification challenges and comparably, or more accurately, classified the samples in the varied datasets.

## Introduction

Single-cell proteomics (SCP) is an expanding field with unique opportunities. SCP is able to measure thousands of proteins per cell, providing researchers the ability to preserve sample heterogeneity at requisite proteomic depth for biological investigations. Because bulk measurements are sample averages, there are several types of biological investigations that rely on single-cell measurements for accurate and meaningful results, including characterizing cell types, identifying changing proteins over the course of development, and mapping a microenvironment and spatial changes at a granular level^1^. With the focus of recent advances in the field shifting toward increasing sample throughput, biological investigation at scale in single-cell proteomics will soon be possible^2–4^.

For many single-cell biological investigations, the first step in the analysis is to determine what types of cells are in the sample and apply a label (cell annotation). While there are experiments that begin with labeled cells, most often cells are unlabeled when measured due to the nature of sample preparation. There are several methods for determining the identity of a cell within an experiment, including external labeling (using known biomarkers) and data-derived labeling (using machine learning). In many cases, external labeling is experimentally challenging or unavailable for the cell types within a sample, increasing reliance on data-derived labeling methods.

In our recent analysis of diverse uses of single-cell technologies, we identified two main machine-learning methods used for data-derived labeling in single-cell analyses: clustering and classifiers^1^. Generally, in cell annotation through clustering, gene/protein abundance measurements are used to group samples together based on the similarity of their abundance profiles (for example, k-nearest neighbors or leiden clustering), and labels are applied by comparing cluster features with lists of canonical markers for the anticipated cell types in the sample^5–7^. Alternatively, in cell annotation through classifiers, a machine learning model is typically trained on the abundances in a reference dataset containing the cell types of interest. This trained model is then used to predict the labels for cells in the unlabeled, experimental dataset^7–9^.

These annotation methods (as currently implemented) come with several challenges that drastically reduce the practicality of using these methods in biological investigations: (1) double dipping, (2) missing-value imputation, and (3) batch effects. Data-derived labeling methods based on clustering or classifiers typically use abundance measurements as features for clustering or classification. Once these measurements are used to group and/or label the cells, using them again in downstream analyses, such as differential expression or developmental trajectory analyses, results in artificially inflated significance and invalid results due to the circular analysis^10–13^. Though a known problem, this pattern of double dipping is baked into published pipelines commonly used in single-cell investigations^5–7^. Furthermore, in classifiers, double dipping can occur when the same data is used to generate and validate the model^14,15^. Beyond issues of double dipping, many traditional clustering and classification algorithms and implementations cannot natively handle null (missing) values in the input data, resulting in missing-value imputation as a common pre-processing step. Missing data is a known problem in proteomics, with null values caused by both non-random effects (below the limit of detection) and random effects (stochasticity of LC-MS data acquisition and analysis)^16,17^. With lower protein volumes in SCP as compared to bulk measurements, a larger proportion of data points are imputed^18^. Because missing values in SCP come from a variety of sources (including low abundance, measurement stochasticity, biological condition, etc.), accurately replacing null values is difficult, and can drastically change downstream results and obscure true biological variation^18^. With the emergence of methods for differential expression on unimputed data^19^, removing imputation as a necessary step in classification would allow for complete investigative workflows that natively handle missing values.

Finally, both labeling methods assume that protein measurements are directly comparable across all samples (within and between the reference and experimental datasets). Batch effects introduced during sample preparation, data acquisition, and data processing all reduce the comparability of protein measurements, and thereby annotation accuracy^20^.

Here we present NIFty, a classifier-based cell annotation tool that performs feature selection, model training and validation, and model application. Using features generated following the principles from top-scoring pairs (TSP)^21,22^, NIFty is the first classification implementation with proteomics in mind that addresses all three challenges associated with data-derived cell annotation: double dipping, missing-value imputation, and batch effects. NIFty’s required input is protein quantitation information (for any number of proteins and generated by any search tool) and class labels (for reference samples) for each sample. Using a variety of SCP datasets, we tested NIFty’s ability to select features and classify samples on unimputed data compared to imputed data, datasets with large batch effects, and a multiclass dataset. In each case, NIFty was successfully able to overcome challenges associated with missing values and batch effects, performing comparably, or better, on the unimputed and non-batch-corrected versions of the tested datasets.

## Results

### NIFty Design

NIFty is a classifier-based cell annotation tool that can be used for feature selection, classification model generation, and/or model application. The code and documentation for NIFty can be found at https://github.com/PayneLab/nifty.

NIFty’s core innovation is in how features are generated, selected, and evaluated for use in machine learning model training and application. Traditionally, the features used for classification of proteomics data are protein abundance values. There are several challenges associated with using abundance measurements as features: (1) datasets with deep proteome coverage result in an imbalanced sample-to-feature ratio in model training; (2) comparing the abundance measurements between samples in the annotation stage precludes their use in downstream analyses (e.g. differential expression); (3) missing values must be imputed before many classifiers can be trained on or applied to the data; and (4) the abundance values between all reference and experimental datasets must be directly comparable (e.g. batch effect corrected and/or normalized).

To overcome these four challenges, NIFty uses an improved implementation of the top-scoring pairs (TSP) concept^21,22^. The features in NIFty are pairwise protein comparisons within samples, rather than comparisons of a single protein’s abundance between samples (see Figure 1A). Each feature is a rule comparing two proteins. For example, a feature/rule might be “protein 1 is greater than protein 2” (protein 1 > protein 2). When evaluating this rule in a sample, if the statement is true, the feature matrix would contain 1; if the statement is false, the matrix would contain 0. The feature matrix is generated by evaluating rules for every combination of two proteins. By comparing proteins within samples, protein abundances do not need to be comparable between samples for accurate annotation, overcoming challenges with batch effects. Additionally, this comparison of protein pairs within samples means that proteins can be compared between samples for the first time in downstream differential expression analyses, avoiding problems of double dipping.

**Figure 1:**
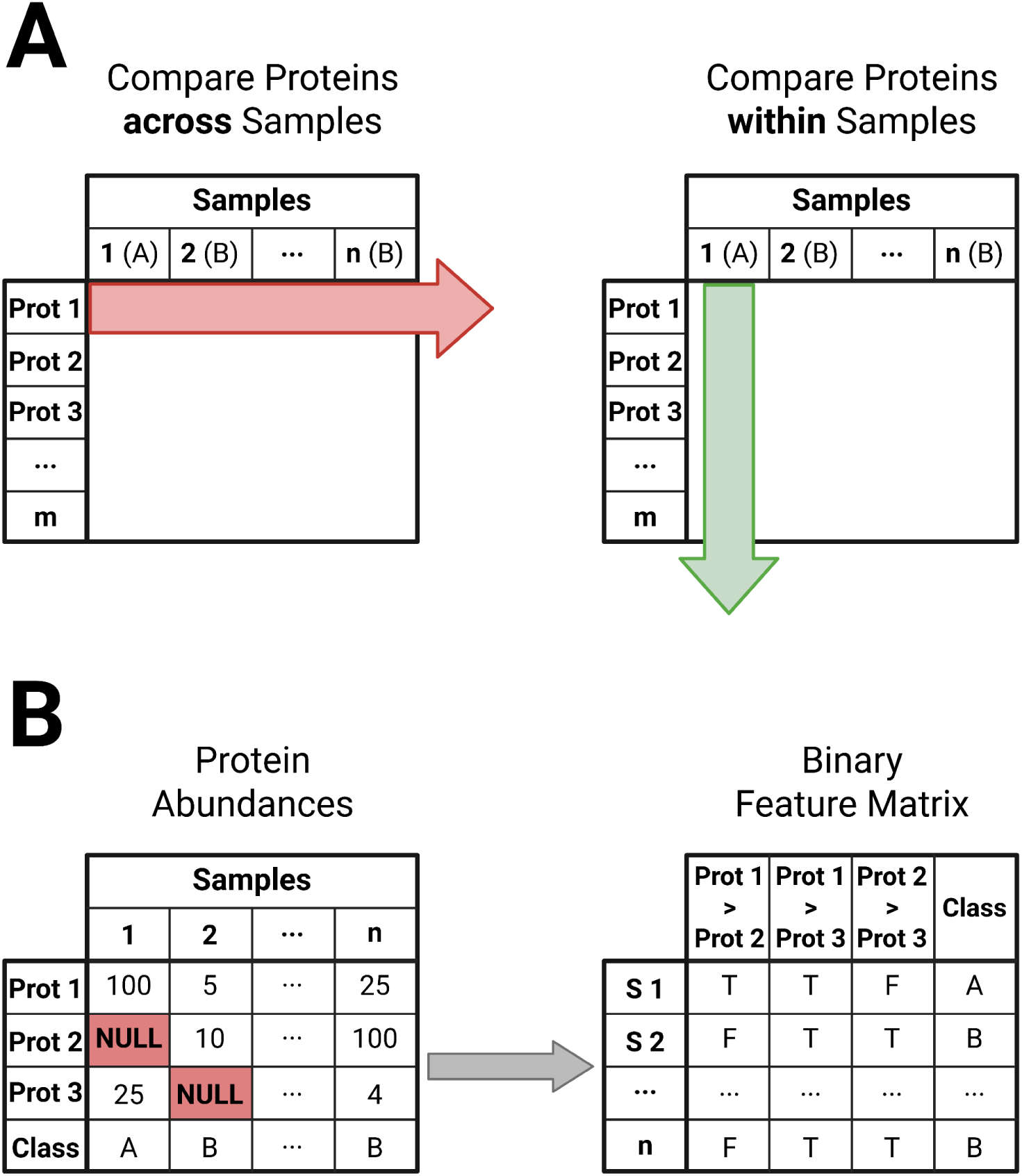
Generating machine-learning features through top-scoring pairs. Most machine-learning classification implementations use protein abundances as features and compare those abundances across samples (**A, left**). This leads to issues with double dipping as abundance measurements are compared in classification and then compared <u>a second time</u> in differential expression or other downstream analyses. Additionally, in these methods, abundance measurements must be comparable across all samples within an experiment, something not easily achieved in proteomics due to large batch effects from multiple sources. In top-scoring pairs, protein abundances are compared within a sample (**A, right**), in the form of pairwise rules (e.g. protein 1 > protein 2) to create a binary feature matrix. By confining abundance comparisons to a sample, batch effects affecting the comparability of measurements between samples are no longer problematic. Additionally, by allowing a rule comparison to be both “protein 1 is greater than protein 2” or “protein 1 is present and protein 2 is absent”, a complete feature matrix can be generated from an incomplete data matrix, overcoming the need to impute missing abundance values in data pre-processing (**B**).

Proteomics data, especially single cell proteomics data, has many missing values in the quantitative data table. These missing values can have several causes, including missing due to low abundance and missing at random^17^. Traditional machine learning software tools require complete data. Therefore, these missing values are typically imputed before classification. Even in other implementations of the TSP method, missing values are imputed prior to rule generation^23,24^. However, NIFty can tolerate missing data by reimagining a rule as being one of two options: “protein 1 is greater than protein 2”, or “protein 1 is present and protein 2 is absent” (see Figure 1B). Thus, a partial quantitative data matrix can still be used to generate a complete feature matrix, eliminating the need for missing-value imputation in pre-processing. NIFty does not make any assumptions about whether the data is missing at random or not. It simply allows presence/absence to be evaluated as part of the rule.

NIFty’s implementation of rules as features solves the issues of double dipping and missing values, and some challenges associated with batch effects. However, they do not solve the issue of too many features relative to the number of samples. In general, machine learning performs well when the number of samples is much greater than the number of features. Without any down-selection, the TSP method generates 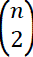 features from *n* proteins – an explosive worsening of the problem. In an SCP dataset with 1,000 proteins, the TSP method creates ∼500,000 rules.

#### Feature Selection

Identifying which features to use in model training is a central part of machine learning, typically called ‘feature selection’. In the TSP paradigm, features are the rules that compare two proteins. Hundreds-of-thousands to millions of rules are created and only a very limited number can be effectively used in model training. Rules need to be scored, sorted, and selected in a computationally efficient manner. Additionally, scoring metrics need to be granular enough to meaningfully separate rules from each other so that the *k* rules selected are the best features for classification.

Previous implementations of TSP do a version of feature selection by down-selecting the proteins (for example, differential expression or random forest feature importance) before generating rules on the new subset to reduce the number of rules that need to be scored and filtered^23,24^. Then, rules are scored and the top *k* rules are output and used to train classifiers. This method of reducing the feature space compares the protein abundances between classes upfront, a form of double dipping. Additionally, this form of feature selection may eliminate proteins that, while not differentially expressed between classes, reliably evaluate in pairs and would have generated high-scoring features.

In contrast, NIFty does not use differential expression to help with feature selection; instead it has simple and computationally robust methods to score, sort and select features. First, to reduce the number of proteins used to generate rules, NIFty implements a filter for the proportion of missing values. For each protein, if the proportion of missing values in all classes is above an acceptable missingness threshold, the protein is dropped. Next, NIFty generates rules from all combinations of the remaining proteins. Once rules have been generated, it is important to assess how well a rule might perform as a feature in a classifier, so NIFty scores each rule based on how characteristic it is of a single class (see Figure 2)^21^.

**Figure 2:**
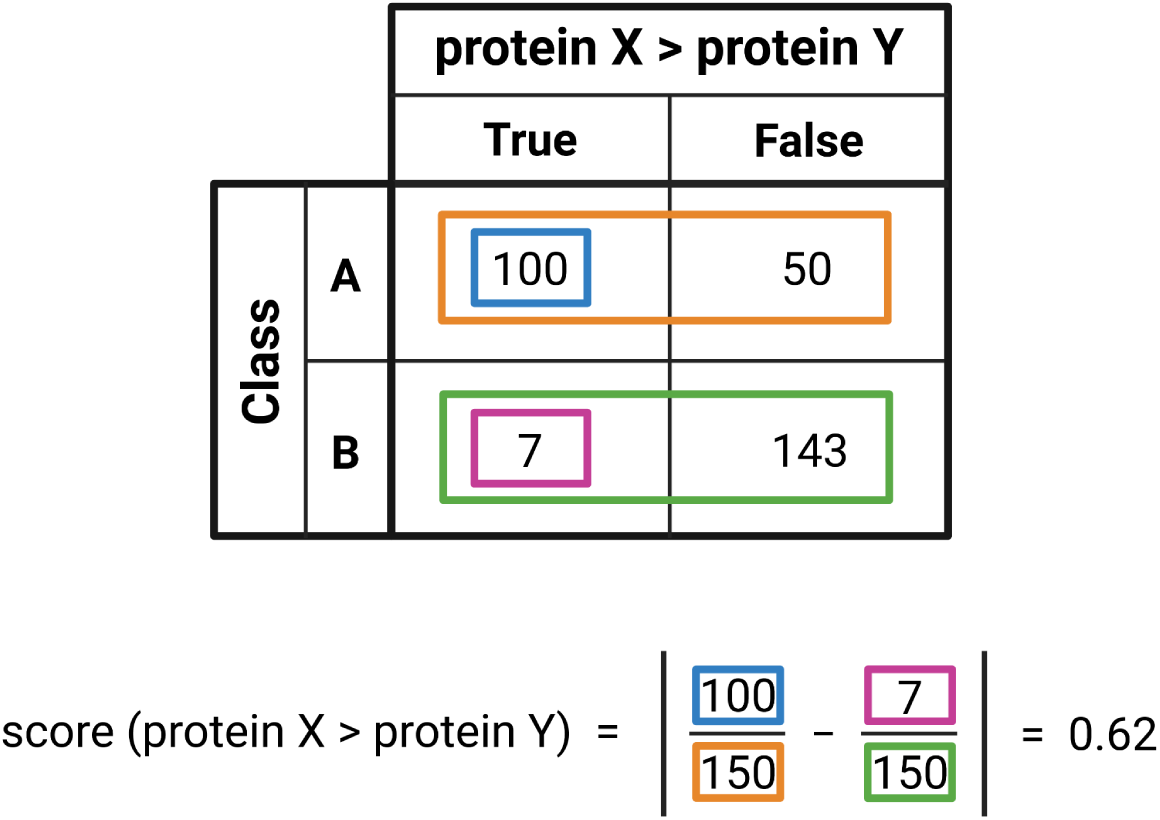
Scoring TSP features. For each rule, a score is generated based on the proportion of ‘true’ hits in one of the classes minus the proportion of ‘true’ hits in the other class. The absolute value is taken to ensure that no matter which class a rule is characteristic of, high scores indicate a rule likely to perform well as a feature in classification. Scores range between zero (no association with either class) and one (perfect association with one of the classes).

To determine the significance of each score and add another layer of granularity to rule rankings, NIFty uses permutation tests to generate a p-value for the score of each rule. Most often, generating a meaningful null distribution and p-value requires the dataset to be randomized at least 100 times and scores generated for each of these permutations. However, because the NIFty feature matrix is binary, rules with similar proportions of ‘true’ and ‘false’ values are already random permutations of each other. This allows us to bin rules together based on the proportion of ‘true’ and ‘false’ values. Labels are randomized and rules re-scored once, as scores from the randomization make up the null distributions for each bin of similar rules (see Methods). Finally, true scores for each rule can be compared to the matching null distribution to calculate a p-value in a more computationally efficient manner than traditional permutation methods. Rules are then ranked based on the lowest p-values and highest scores.

The final aspect used in selecting rules is whether a high-scoring (and low p-value) rule provides a unique set of information. To ensure that the top *k* features selected present the most information for the machine learning classification, a mutual information score is used to filter rules out. If a top-ranking rule shares high amounts of mutual information with at least one other rule that has already been selected, the rule is not selected and lower ranking rules are considered (see Methods). Giving up some redundant, high-scoring rules in favor of rules that provide new information about the classes benefits classification in the long run. The final set of selected features is then used to train a machine learning classifier for use on unlabeled experimental data.

### NIFty Performance

To demonstrate NIFty’s utility in a variety of situations, we report NIFty’s performance on cell type classification using imputed vs. unimputed data, data with large batch effects, and a multiclass dataset. Overall, NIFty performed well in all categories.

#### Testing on Incomplete Data

We first wanted to ensure that NIFty performed comparably on imputed and unimputed data, validating NIFty’s method of tolerating missing values in the input. To investigate this, we used NIFty to classify cell type with datasets from Leduc et al.^25^ and Montalvo et al.^26^. The dataset from Leduc et al. has cell types that are easily distinguishable, while the dataset from Montalvo et al. has cell types that are more difficult to distinguish. We ran 100 trials at eight different input sizes on both the imputed and unimputed data for each experiment. We randomly selected samples in each trial, ranging from 25 to 300 samples per class, to be used in feature selection (30% of the samples) and model training (70% of the samples). We randomly selected and held out 50 samples per class as a validation set for every trial. With 50 samples per class or more, on average NIFty performed comparably, or slightly better, on unimputed data than on imputed data (see Figure 3).

**Figure 3:**
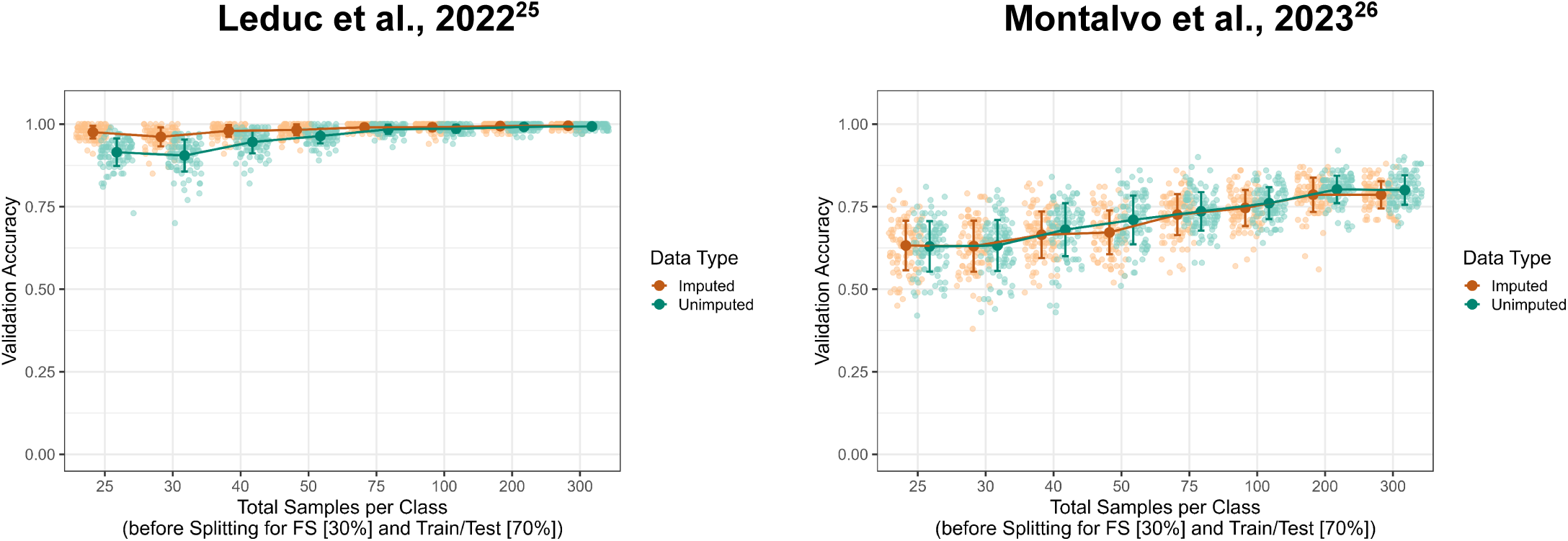
NIFty classification accuracy comparing imputed and unimputed data. Each dataset was run across a wide variety of input sizes, on both an imputed (orange data points) and unimputed (teal data points) quantification matrix. Each light data point represents the validation accuracy for one trial. Each dark datapoint represents the average validation accuracy from 100 trials. Error bars represent +/− one standard deviation. For each trial, between 25 and 300 samples per class were randomly selected. These samples were then split between feature selection (30% of the samples) and model training/testing (70% of the samples). An additional 50 samples per class were held out as a validation set for every trial. Importantly, the average validation accuracy between the imputed and unimputed data for trials with 50 or more total samples per class were nearly identical.

We then tested NIFty on imputed and unimputed data from a wider variety of datasets^27–31^. These datasets were generated by multiple labs, on different LC-MS instruments, with different sample preparation and acquisition methods, and represent both primary cells and cell lines. We ran 100 trials at two different input sizes on both the imputed and unimputed data for each experiment. We randomly selected 50 or 100 samples per class in each trial to be used in feature selection (30% of the samples) and model training (70% of the samples). We randomly selected and held out 50 samples per class as a validation set for every trial. The results of these tests can be found in Table 1. The median difference between the average imputed validation accuracy and average unimputed validation accuracy was 0.6% for tests run with 50 samples per class and 0.3% for tests run with 100 samples per class.

**Table 1:** NIFty classification accuracy comparing imputed and unimputed data. Each dataset was run with two input sizes on both an imputed and unimputed quantification matrix. The results reported for Leduc et al. and Montalvo et al. match those reported in Figure 2 for 50 and 100 total samples per class. Each reported accuracy in the table is an average validation accuracy from 100 trials. For each trial, either 50 or 100 samples per class were randomly selected. These samples were then split between feature selection (30% of the samples) and model training/testing (70% of the samples). An additional 50 samples per class were held out as a validation set for every trial. In most cases, the average accuracy between the imputed and unimputed data was nearly identical. We note that two in two cases, the reported accuracy for the unimputed data was noticeably lower than the reported accuracy for the imputed data.

| Dataset<br>(PMID) | Cell Types | Validation Accuracy<br>50 Samples / Class |  | Validation Accuracy<br>100 Samples / Class |  |
| --- | --- | --- | --- | --- | --- |
|  |  | Imputed | Original | Imputed | Original |
| Ai et al.<br>(39855627) | iPSC day0<br>iPSC day21 | 0.9587 | 0.9720 | 0.9748 | 0.9886 |
| Furtwängler et al.<br>(40839704) | HSC<br>EMP | 0.8069 | 0.7315 | 0.8288 | 0.7698 |
| Furtwängler et al.<br>(40839704) | HSC<br>Early Eryth | 0.9664 | 0.9602 | 0.9860 | 0.9718 |
| Khan et al.<br>(39416047) | SPG<br>St | 0.9009 | 0.8895 | 0.9198 | 0.9129 |
| Leduc et al.<br>(36527135) | WM989<br>U937 | 0.9823 | 0.9638 | 0.9906 | 0.9859 |
| Montalvo et al.<br>(37609178) | Islet β-cell Dec1 KO<br>Islet β-cell WT | 0.6720 | 0.7098 | 0.7459 | 0.7606 |
| Petrosius et al.<br>(40334743) | HEK293<br>U937 | 0.9670 | 0.9829 | 0.9819 | 0.9912 |
| Saddic et al.<br>(40027651) | MFN smc<br>WT smc | 0.5987 | 0.5844 | 0.6750 | 0.6129 |
| Saddic et al.<br>(40027651) | MFN fibro<br>WT fibro | 0.6698 | 0.6447 | 0.7168 | 0.7135 |
| Saddic et al.<br>(40027651) | MFN smc<br>MFN fibro | 0.9893 | 0.9844 | 0.9925 | 0.9924 |
| Saddic et al.<br>(40027651) | WT smc<br>WT fibro | 0.9855 | 0.9851 | 0.9885 | 0.9932 |

NIFty’s ability to tolerate incomplete (unimputed) data is critical when using data from a different experiment as a reference or combining multiple datasets into a reference (e.g. cell atlases). Without the ability to handle missing information, all missing values would either need to be imputed or the datasets would need to be filtered down to complete proteins (proteins with no missing values). Within the experiments in the above table, 3% (+/− 2.5%) of the proteins had no missing values in the combined reference dataset (feature selection + training + validation) among all trials (median +/− IQR). Additionally, 0% (+/− 13.3%) of the top 15 rules selected per trial had no missing values. The likelihood of having complete proteins between a reference and experimental dataset, or after aggregating multiple datasets, is near-zero due to measurement stochasticity and both experimental and analytical batch effects. Even in large QC datasets where the exact same sample is repeatedly analyzed, zero proteins were identified in every sample^32^.

#### Testing on Data with Large Batch Effects

Next, we evaluated NIFty’s performance on data containing large, uncorrected batch effects. The features generated by NIFty for machine learning always compare protein measurements within samples, and never between samples, resulting in comparisons impervious to issues accompanying uncorrected batch effects.

In a classification task, batch effects are at their worst when one dataset (or batch) is used to generate a model that is then applied to a completely separate dataset. The more datasets, or batches, included in the generation of the model, the better the model will generalize when applied to new data. With this in mind, we tested NIFty on a dataset created by the HUPO Single Cell Initiative containing single-cell samples of two different cell types measured in eight different batches. Each batch consisted of 105-108 cells of each cell type, and contained batch effects due to a combination of differing data acquisition methods and instruments, and data processing batches. Two quantification tables were generated from the raw data: one that combined samples from all batches and performed cross-run normalization, and one that combined samples from all batches but did not include cross-run normalization. We ran NIFty on both the normalized and non-normalized quantification tables. In each trial of NIFty on the data, one batch was held out as a validation set, while at least one other batch was used in model generation. For control trials, samples were randomly selected from one batch for both model generation and model validation to act as a baseline maximum for validation accuracy. We ran 100 trials for each possible combination of batches. Samples from batches selected for model generation were pooled and we randomly selected 50 samples per class per batch in each trial to be used in feature selection (30% of the samples) and model training (70% of the samples). We randomly selected and held out 50 samples per class from the validation batch as a validation set for every trial. The results of these tests can be found in Figure 4. With three or fewer batches used in training, classification accuracy was better using non-normalized data. For larger numbers of batches, NIFty performed indistinguishably on the non-normalized data versus the normalized data in all cases.

**Figure 4:**
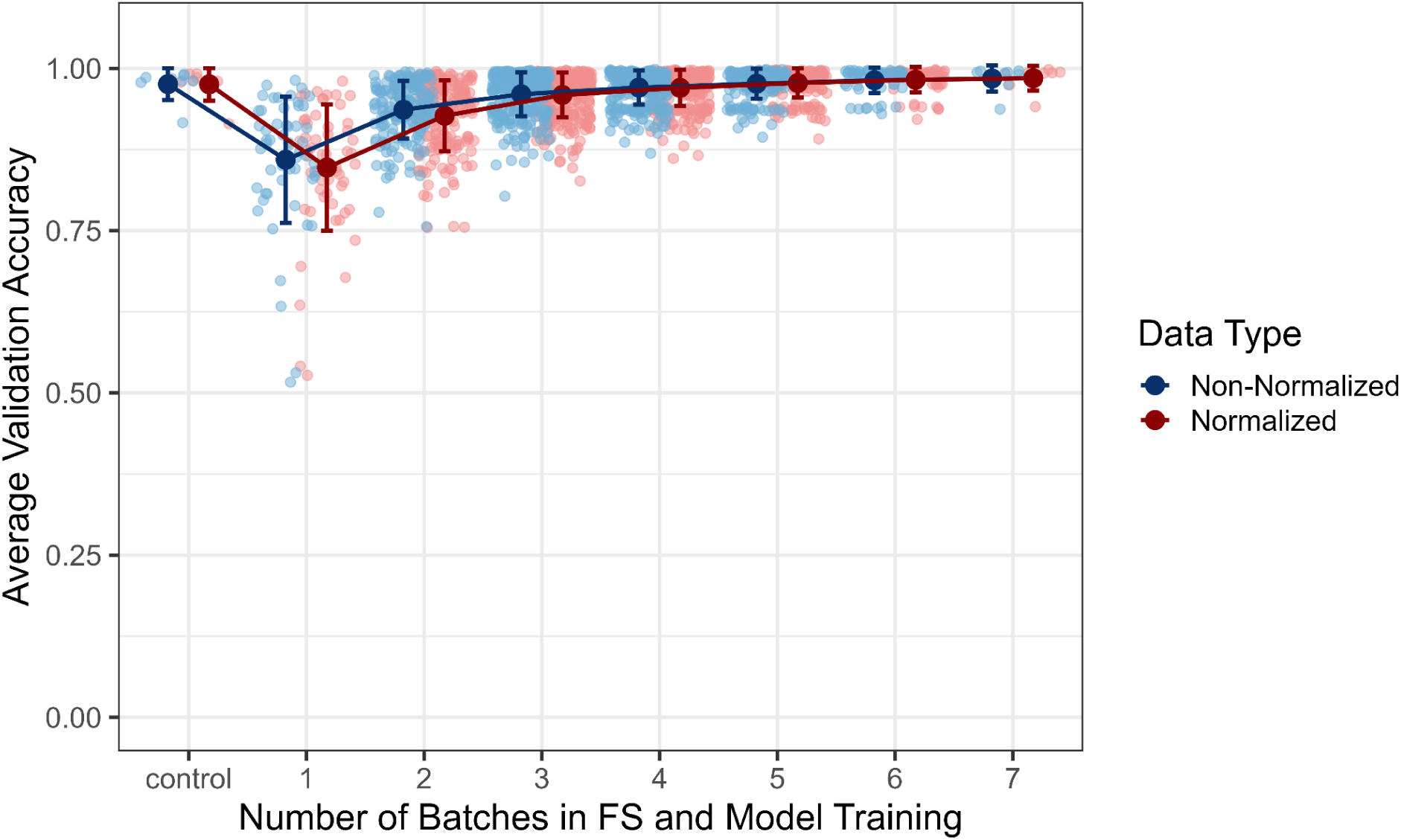
NIFty classification accuracy comparing normalized and non-normalized multi-batch data. Each combination of batches was run on both a normalized (red data points) and non-normalized (blue data points) quantification matrix. In control trials, the same batch was used for both model generation (feature selection and model training) and model validation, representing a baseline maximum for validation accuracy within each batch (without sample overlap between model generation and validation). In all other trials, a varying number of batches were used for model generation, while one batch was held out and used for model validation. Each light data point represents one possible combination of batches and reports the average validation accuracy for 100 trials. Each dark data point represents the average validation accuracy from all trials of all possible combinations of batches. Error bars represent +/− one standard deviation. For each trial, samples from batches selected for model generation were combined. 50 samples per class per batch were randomly selected from the combined samples and then split between feature selection (30% of the samples) and model training/testing (70% of the samples). An additional 50 samples per class were randomly selected and held out for model validation from the batch selected for model validation. In all cases, the overall average validation accuracy for the non-normalized data was comparable to, or slightly better than, that of the normalized data.

#### Testing on Multiclass Data

As a final data classification task, we evaluated NIFty on multiclass data. We anticipate that many of the datasets created for biological investigation in the future will not be datasets consisting of only two classes of cells. To enable multiclass, we first use NIFty to identify features that distinguish a single class in a one-vs-rest scenario. After generating features for each individual class, this larger set of features is used in a multiclass classification method in Scikit-learn^33^ (see Figure 5).

**Figure 5:**
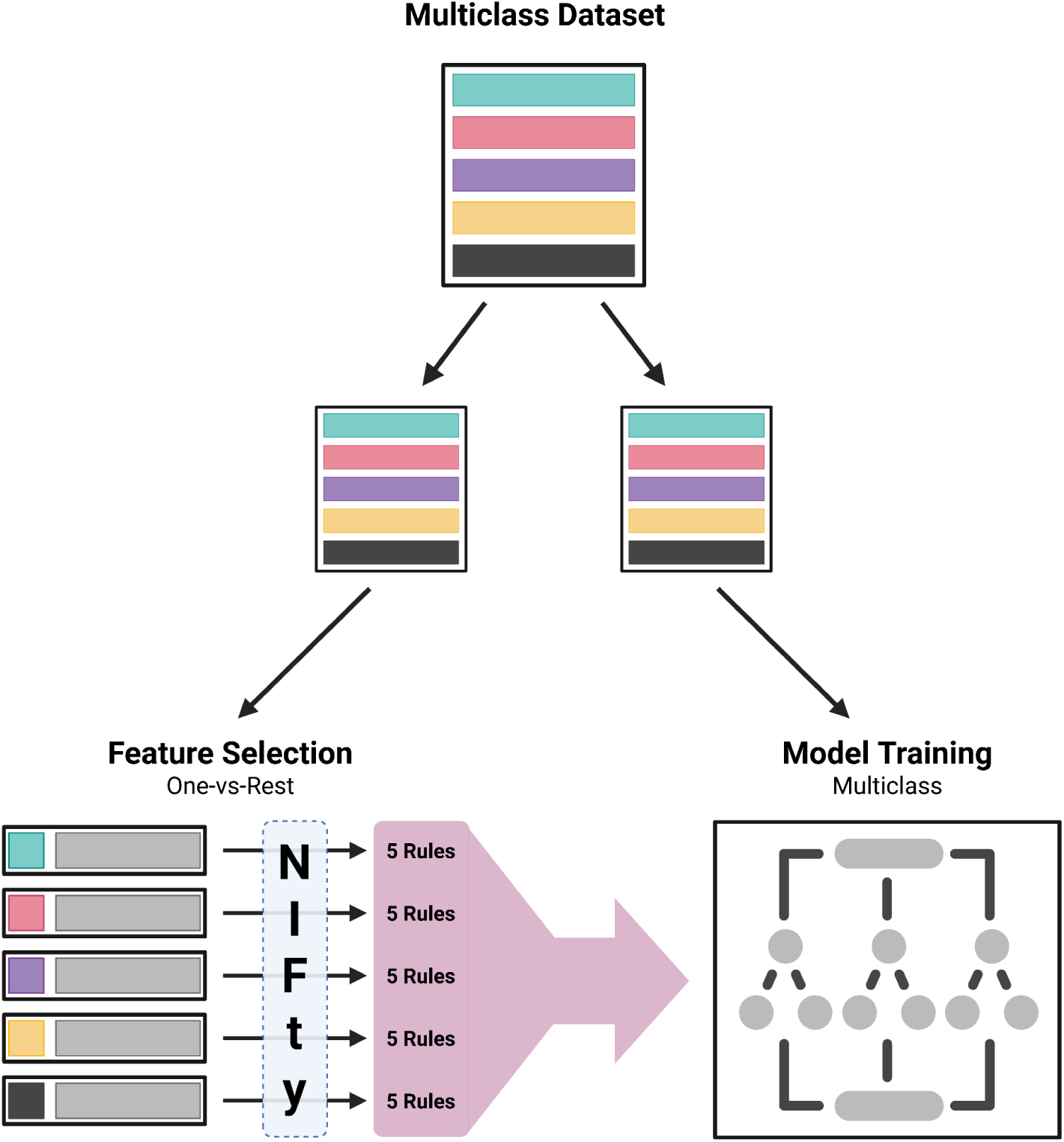
Using NIFty on multiclass data. A multiclass reference dataset is split into several non-overlapping sets, a feature selection set and a model training set (which is further split into a training set and a validation set). To generate features, the feature selection set is run through NIFty once for every class in a one-vs-rest scheme. In this scheme, each class retains its true label in one run and is run against all other classes in an “other” group. In this way, NIFty is able to generate a set of features characteristic to an individual class in each run. These features are aggregated and used as the complete feature set in training a multiclass machine learning model. This model can then be used to predict labels for multiclass experimental datasets.

Using this framework, we ran NIFty on the data from Ai et al., where human induced pluripotent stem cell (iPSC) cells were sampled and measured at various timepoints in their development towards cardiomyocytes (days 0, 2, 4, 10, and 21)^27^. We used the different timepoints as classes and ran NIFty five separate times on a portion of the data in a one-vs-rest scheme (day 0 vs others, day 2 vs others, etc.). Five rules/features were selected each time NIFty was run, resulting in 25 unique rules used as features in model training. We ran 100 trials of this experiment. In each trial, 50 samples per class were randomly selected for the feature selection dataset, 100 samples per class were randomly selected for the model training dataset, and 50 samples per class were held out and used as a validation set. For most stages of cell development, NIFty was highly accurate in its ability to separate classes. The final two timepoints, day 10 and 21, are both considered cardiomyocytes (“non-purified contracting iCMs” and “differentiated purified contracting iCMs”, respectively)^27^. Classifying day 10 cells was the most difficult, with 86% accuracy and errors largely being a misclassification as day 21 cells (see Figure 6).

**Figure 6:**
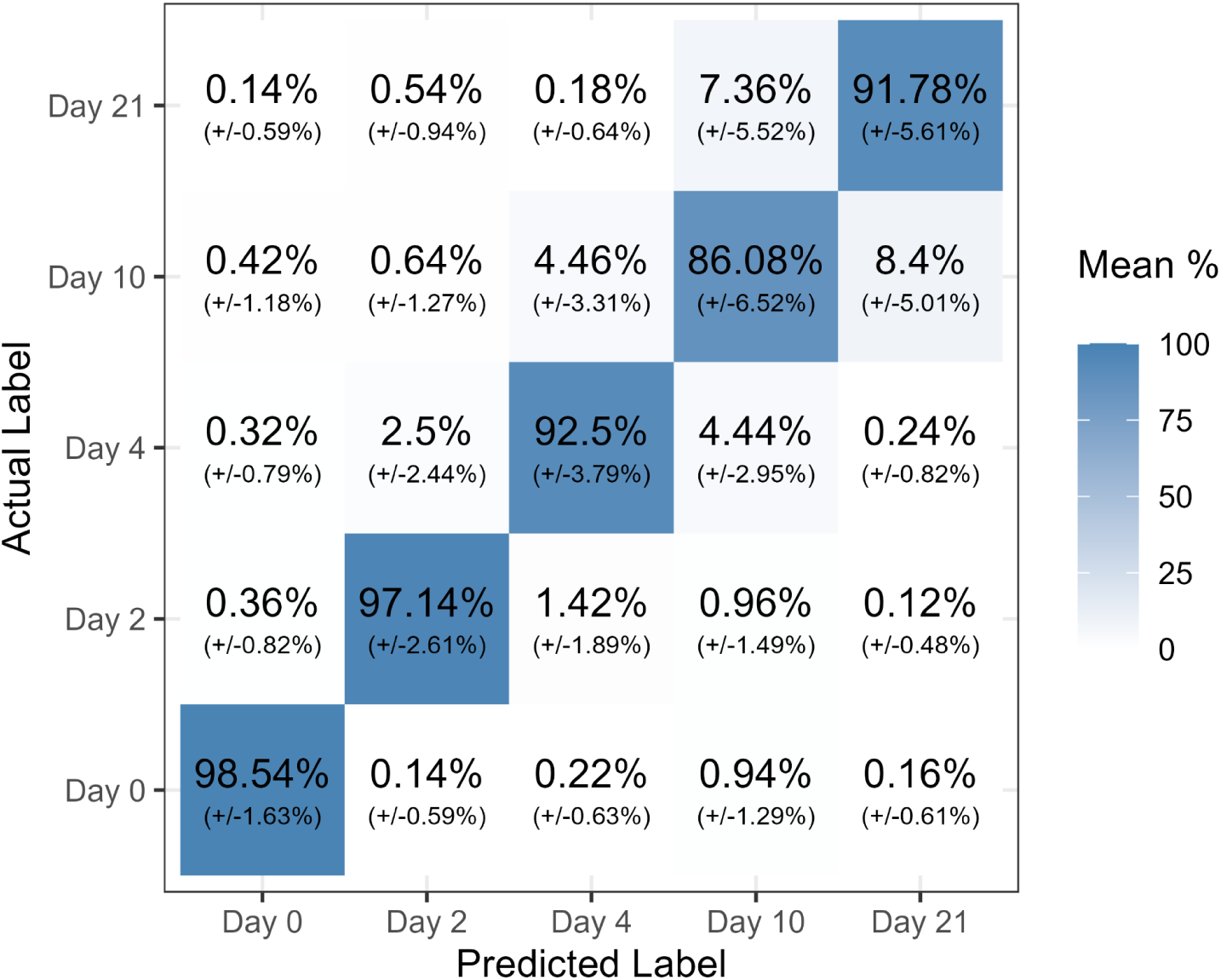
Classification accuracy when testing NIFty on a multiclass dataset. Each tile represents the average percentage of validation-set observations (+/− one standard deviation) that fell into that combination of actual and predicted labels from 100 trials. For each trial, 50 samples per class were randomly selected for feature selection and 100 samples per class were randomly selected for model training. Additionally, 50 samples per class were randomly selected and held out as a validation set for each trial. In most cases, the actual and predicted labels matched (dark band along the diagonal). We note that both Day 10 and Day 21 represent cardiomyocytes^27^. Accurately classifying Day 10 cells was the most difficult, with most misclassifications being Day 21.

## Discussion/Conclusion

We present the NIFty software tool for cell labeling and classification, and demonstrate its performance in a variety of classification scenarios for single-cell proteomics. By adapting the top-scoring pairs concept to explicitly handle missing data and fixing the feature selection, our tool overcomes three critical problems associated with cell-labeling algorithms and tools. First, NIFty does not require complete data and therefore does not force researchers to impute. Second, NIFty does not compare protein abundance across samples during feature selection or training, thus avoiding the pervasive problems with double dipping. Finally, by comparing within samples, NIFty overcomes batch effects. These three critical advantages enable NIFty to facilitate a more effective and straightforward use of single-cell atlases.

The most common use for an atlas is to serve as a hub of integration and a reference of proteome expression for specific cell types. With advances in sample throughput, single cell proteomics will very soon be ready to begin assembling an atlas. The primary benefit to our community of an SCP atlas is that it aggregates a larger reference dataset than a single lab could create on its own. Thus, researchers no longer have to create their own reference dataset to describe baseline proteome expression of a cell type. Additionally, the much larger number of reference cells contained in an atlas is essential in creating a robust model to classify samples. The aggregate of experimental data across many labs and biological replicates within each cell type provides the volume of information necessary for the model to differentiate wanted biological variation from unwanted technical variation. This allows better generalization when transferring labels from a reference to experimental samples, resulting in more accurate annotations and downstream results.

As we anticipate this atlas being created, it is essential that associated algorithms enabling the use of the atlas are developed and vetted. As most SCP data is not generated with an external label, the vast majority of atlas use cases involve data-derived labeling and classification. NIFty represents a different approach to cell classification, one that avoids significant computational and statistical challenges.

The practicalities of atlas creation for single-cell proteomics include a few specific challenges due to the nature of proteomics data creation and processing. Some of the data generation and analysis methods common in our community may make the data incongruent with an atlas.

The first challenge arises when a biological variable overlaps with a technical variable. One example of this can be found in the way that a two-pass search works in most popular DIA tools. If data analysis is batched according to biological variables, there will be many proteins whose presence/absence is perfectly aligned with the biological variable. We saw this in some of the public data that we re-analyzed for this manuscript. Single-cell proteomics often creates hundreds or thousands of data sets, and analyzing these all at one time can be excessively time-intense or impossible with certain software tools. Therefore, some projects chose to perform protein identification/quantification in non-randomized batches. This is problematic as a two pass search (misleadingly called ‘match between runs’ in some DIA software) can lead to significant batch effects in which proteins are reported.

The potential for analytical batches to be created by search tools emphasizes the necessity that an atlas contain data from multiple labs. If a single cell type is only represented by one lab, we can expect annotation accuracy to be significantly impacted (see Figure 4). However, as multiple labs and multiple experiments contribute to an atlas, the features that typify a cell type are generalized and become robust. Based on our analysis of the multi-batch data, we believe that at least three different labs should contribute data describing a cell type for the atlas to be robust.

Another challenging data type is that generated using the popular TMT, or isobaric, multiplexing reagents. These are attractive because of the opportunity to both increase signal to noise in the ultra-low signal space of single-cell proteomics and reduce instrument time. To understand whether these samples can be a productive part of an atlas, we consider the primary purpose of an atlas – to provide reference data about what proteins are expressed in various cell types. By this definition, the data in an atlas must be able to relate to other datasets. Unfortunately, TMT data is normalized across samples by design (i.e. using bridge samples to normalized protein values across plexes). If one were using a cross-sample classifier (which we don’t suggest because of obvious double dipping), then these samples could only serve as a reference if the new data also used the same bridge samples. TMT data is also problematic as a reference for ‘within sample’ feature algorithms like NIFty, because the protein data has been normalized to the bridge sample and therefore comparisons are no longer ‘is protein 1 > protein 2’, but rather ‘is the ratio of protein 1 to a bridge sample > the ratio of protein 2 to a bridge sample’. Here again, the inclusion of a bridge sample in the abundance values means that it cannot serve as a meaningful reference to samples that do not have access to this exact bridge sample. Therefore, we do not believe that TMT data can be part of a single-cell proteomics atlas.

## Methods

Due to our belief that data imputation is unnecessary, NIFty stands for: Never Impute Features, thank you. Code and instructions for running NIFty on your own data can be found here: https://github.com/PayneLab/nifty. Code and instructions for recreating the results presented in this manuscript can be found here: https://github.com/PayneLab/nifty-manuscript. All data and code used to generate figures and tables within this manuscript can be found at https://github.com/PayneLab/nifty-manuscript/tree/main/Figures_and_Tables.

### NIFty Design

NIFty can perform all, or a selection of, three tasks: top-scoring pairs (TSP) rule generation and feature selection, classification model generation, and classification model application.

Feature selection and model generation tasks require reference data containing protein quantitation and metadata information for cells falling into two classes. If one reference dataset is provided for both feature selection and model generation, the dataset is split into three, non-overlapping parts: a feature selection set (15%), a training set (65%), and a validation set (20%). If feature selection is the only task selected, the entire reference dataset is used in this task. If model generation is the only task selected, the dataset is split into two, non-overlapping parts: a training set (70%) and a validation set (30%). If there are an uneven number of samples per class in the validation set, the classes are balanced. A minimum of 15, 35, and 15 samples per class are required for feature selection, model training, and model validation respectively. When run, NIFty generates files containing selected features, trained model, model information, and predicted classes.

#### Rule Generation and Feature Selection

To generate all possible protein-pair rules, NIFty first filters out proteins with more missing values than a threshold value (default 50%) in each class. Next, all combinations of protein pairs are generated to create rules (protein 1 > protein 2). The rules are evaluated to ‘true’ (1) or ‘false’ (0) on the reference data to create a binary feature matrix.

To filter down to the *k* best rules for model training, each rule is scored based on how characteristic it is to one of the two classes. The scoring function used in NIFty is the original scoring function proposed by Gemen et al.^21^, as seen in Equation 1. Once rules are scored, p-values are generated for each score using permutation testing. Rules in the binary feature matrix are binned into 100 groups based on the proportion of ‘true’ and ‘false’ values they contain (rules with similar proportions are binned together). Sample labels are randomized, and all rules are rescored. Null distributions for each bin are generated using the new scores. In cases where fewer than 100 samples fall into a bin, sample labels are randomized again and rules within the bin are rescored, until at least 100 scores make up the null distribution. A p-value for each rule is generated by comparing the observed score for a rule to the null distribution associated with the bin the rule falls into. The formula for our p-value calculations can be found in Equation 2. No p-value correction methods are applied, as p-values are not used for significance in NIFty, but rather as a second score for rule ranking.

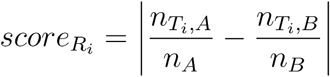

**Equation 1: Rule scoring function.** Given a rule *R_i_*, where *nT_i_,A* is the number of times *R_i_* evaluates to True in class A, *nT_i_,B* is the number of times *R_i_* evaluates to True in class B, *n_A_* is the number of samples in class A, and *n_B_* is the number of samples in class B, the score is calculated by subtracting the proportion of times the rule evaluates to ‘true’ in class B from the proportion of times the rule evaluates to ‘true’ in class A and taking the absolute value of the result.

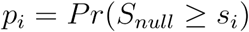

**Equation 2: P-value calculation.** Given a set of possible scores (null distribution), *p_i_* is the probability of observing a score (*S_null_*) at least as large as the observed score (*S_i_*).

Scores and p-values are used to rank the rules (lowest p-values and highest scores). If specified by the user, the rules can be filtered based on a mutual information threshold (default enabled) or a disjoint flag (default disabled). When mutual information filtering is enabled, the top *k* rules that all share less mutual information than a specified threshold value (default 0.7) with each other are selected. When disjoint filtering is enabled, the top *k* rules that do not share any proteins are selected.

A maximum of 50 rules can be selected.

#### Model Generation and Application

Using the selected rules and the training and validation sets, NIFty trains either an SVM or random forest (RF) machine learning model using a stratified k-fold cross validation (CV) strategy (default 5-fold) with Scikit-learn^33^. If there are proteins in the selected rules that are not present in the the training or validation sets, those proteins are be added to the datasets before model generation (quantification filled in with missing values) or the rules containing those proteins are not used as features in model generation (default proteins are added to the training and validation datasets). NIFty uses Scikit-learn’s default hyperparameters when training the classification model. Alternatively, if specified by the user, the models can be trained with either randomized search or grid search hyperparameter tuning.

NIFty reports the following information for the trained model: model hyperparameters; mean and standard deviation for CV accuracy, precision, and recall; and final accuracy, precision, and recall scores based on testing on the held-out validation set.

Once the model is trained, NIFty can use it to predict the class or estimated class probabilities for unlabeled, experimental samples (default class prediction).

### Collecting and Preparing Datasets for Testing

Input data to NIFty is generally two tables: a metadata table which annotates samples, and a quantitative data table of proteome abundances. All datasets required the metadata and quantification data to be formatted in NIFty’s specific input format (see https://github.com/PayneLab/nifty/blob/main/docs/input_file_formats.md). Additional filtering and preparation steps unique to each dataset are described below.

Code to prepare the datasets for NIFty can be found here: https://github.com/PayneLab/nifty-manuscript/tree/main/Data_Prep.

#### Ai et al

`report1.tsv` was downloaded from ftp://massive-ftp.ucsd.edu/v07/MSV000094438/updates/2024-10-23_bineka_5deafa61/other/Supplementary Files/iCMs/

The data was prepared for NIFty by filtering the data to single-cell runs and high quality IDs (`Lib.PG.Q.Value` < 0.01). Protein labels from the `Protein.Group` column and quantification values from the `PG.MaxLFQ` column were used to generate the quantification matrix for NIFty. Proteins with more than 70% missing data in every class were also filtered out. Missing values were imputed using Scikit-learn’s KNNImputer with `n_neighbors` as 3 and `weights` as ‘distance’ to create the imputed dataset. Classification labels for each sample were extracted from the file names.

#### Furtwängler et al

`hBM_scpMS_scRNAseq\results\3_2_integration_validation\hBM_glue_celltyp e.h5ad` was downloaded from https://zenodo.org/records/15554000

The data was prepared for NIFty by extracting the meta, imputed, and unimputed data from the AnnData object. Samples were filtered to the following cell types: ‘HSC’, ‘EMP’, and ‘Early Eryth’. Missing values in the unimputed dataset were filled in with NA values.

#### HUPO Single Cell Initiative

Normalized and non-normalized quantification data was provided by the HUPO Single Cell Initiative, with anonymized cell type and batch ID labels.

The data was prepared for NIFty by filtering the data to high quality IDs (`Global.PG.Q.Value` < 0.01). Protein labels from the `Protein.Group` column and quantification values from the `PG.MaxLFQ` column were used to generate the quantification matrix for NIFty.

#### Khan et al

`npop5_proteinMatrix_Imputed.BCmTRAQ.txt` and `npop5_proteinMatrix_NoImp.BCmTRAQ.txt` were downloaded from https://drive.google.com/drive/folders/1iboBCc1zxTvzfxZM8VoS_EZZrFVDuSlI `Protein_cellTypeLabels_postAlignment.txt` was downloaded from https://drive.google.com/drive/folders/1LGgUilllIlUU8BvM8Tj_IUMsep5_xkeX

The data was prepared for NIFty by filtering the data to runs that were part of experiment five. Samples were filtered to the following cell types: ‘SPG’ and ‘St’.

#### Leduc et al

`t6.csv` (imputed data), `t7.csv` (unimputed data), and `meta.csv` were downloaded from https://drive.google.com/drive/folders/12-H2a1mfSHZUGf8O50Cr0pPZ4zIDjTac

The data was prepared for NIFty by filtering the samples to the following cell types: ‘m’ and ‘u’.

#### Montalvo et al

`Protein_imputed.csv`, `Protein_uniputed.csv`, and `meta.csv` were downloaded from https://drive.google.com/drive/folders/1xHlUsGOfdpdL3byPbcSTX7OYU-dw02_T

The data was prepared for NIFty by filtering the samples to the following cell types: ‘ko’ and ‘wt’.

#### Petrosius et al

Raw data for scHEK samples was downloaded from ftp://massive-ftp.ucsd.edu/v08/MSV000095333/raw/Exp6/scDataset/, ftp://massive-ftp.ucsd.edu/v08/MSV000095333/raw/Exp6/scDataset/pt1/, ftp://massive-ftp.ucsd.edu/v08/MSV000095333/raw/Exp6/scDataset/pt2/, and ftp://massive-ftp.ucsd.edu/v08/MSV000095333/raw/Exp6/scDataset/pt3/

Raw data for CD34 samples was downloaded from ftp://massive-ftp.ucsd.edu/v08/MSV000095333/raw/Exp6/human_CD34/

Raw data for U937 samples was downloaded from ftp://massive-ftp.ucsd.edu/v08/MSV000095333/raw/Exp6/u937/

The raw data was searched with DIA-NN (v 2.2.0 Academia)^34^. A spectral library was generated from a UniProt human proteome download containing reviewed sequences, isoforms, and contaminants (downloaded on 2025-12-09). Using the predicted spectral library and proteome download from UniProt, the raw data was searched with the following changes to default settings: spectral library generation was disabled, mass accuracy was set to 10.0, MS1 accuracy was set to 4.0, MBR was disabled, and cross-run normalisation was set to RT-dependent. This generated the `report.parquet` file used for testing.

The data was prepared for NIFty by filtering the data to scHEK and U937 runs and high quality IDs (`Global.PG.Q.Value` < 0.01). Protein labels from the `Protein.Group` column and quantification values from the `PG.MaxLFQ` column were used to generate the quantification matrix for NIFty. Contaminants and proteins with more than 70% missing data in every class were also filtered out. Missing values were imputed using Scikit-learn’s KNNImputer with `n_neighbors` as 3 and `weights` as ‘distance’ to create the imputed dataset. Classification labels for each sample were extracted from the file names.

#### Saddic et al

Unimputed quantification data and cell-type labels were provided by Saddic et al.

The data was prepared for NIFty by filtering the samples to the following cell types: ‘smc’ and ‘fibro’. Proteins with more than 70% missing data in both classes were also filtered out. Missing values were imputed using Scikit-learn’s KNNImputer with `n_neighbors` as 3 and `weights` as ‘distance’ to create the imputed dataset.

### NIFty Performance Tests

#### Trials and Results

The trials that generated the results presented were batch run through Brigham Young University’s supercomputing resources. For each trial, we randomly selected a subset of the available samples and generated the trial-specific quantification, metadata, and configuration files.

Code for random sampling and input-file generation for testing on incomplete data and result aggregation can be found here: https://github.com/PayneLab/nifty-manuscript/tree/main/Testing/No_Imputation.

Code for random sampling and input-file generation for testing on multiclass data and result aggregation can be found here: https://github.com/PayneLab/nifty-manuscript/tree/main/Testing/Multiclass.

## Acknowledgements

The authors thank the HUPO Single Cell Initiative and Sarah Parker (Cedars-Sinai Medical Center, Los Angeles, CA) for sharing pre-publication data. This work was supported by an NIGMS/NIH award (R01GM147653 and R01GM147653-02S2 to S.H.P. and R01GM147653-02S1 to A.A.N.), with additional support from the BYU Simmons Center for Cancer Research and the BYU College of Life Sciences Undergraduate Research Awards.

## Notes

### Competing Interest Statement

The authors have declared no competing interest.

## References

(1) Nitz, A. A.; Giraldez Chavez, J. H.; Eliason, Z. G.; Payne, S. H. Are We There Yet? Assessing the Readiness of Single-Cell Proteomics to Answer Biological Hypotheses. J. Proteome Res. 2025, 24 (4), 1482–1492. 10.1021/acs.jproteome.4c00091.

(2) Slavov, N. Scaling Up Single-Cell Proteomics. Mol. Cell. Proteomics MCP 2022, 21 (1), 100179. 10.1016/j.mcpro.2021.100179.

(3) Sanchez-Avila, X.; de Oliveira, R. M.; Huang, S.; Wang, C.; Kelly, R. T. Trends in Mass Spectrometry-Based Single-Cell Proteomics. Anal. Chem. 2025, 97 (11), 5893–5907. 10.1021/acs.analchem.5c00661.

(4) Ghosh, G.; Shannon, A. E.; Searle, B. C. Data Acquisition Approaches for Single Cell Proteomics. Proteomics 2025, 25 (1–2), e2400022. 10.1002/pmic.202400022.

(5) Hao, Y.; Stuart, T.; Kowalski, M. H.; Choudhary, S.; Hoffman, P.; Hartman, A.; Srivastava, A.; Molla, G.; Madad, S.; Fernandez-Granda, C.; Satija, R. Dictionary Learning for Integrative, Multimodal and Scalable Single-Cell Analysis. Nat. Biotechnol. 2024, 42 (2), 293–304. 10.1038/s41587-023-01767-y.

(6) Wolf, F. A.; Angerer, P.; Theis, F. J. SCANPY: Large-Scale Single-Cell Gene Expression Data Analysis. Genome Biol. 2018, 19 (1), 15. 10.1186/s13059-017-1382-0.

(7) Heumos, L.; Schaar, A. C.; Lance, C.; Litinetskaya, A.; Drost, F.; Zappia, L.; Lücken, M. D.; Strobl, D. C.; Henao, J.; Curion, F.; Single-cell Best Practices Consortium; Schiller, H. B.; Theis, F. J. Best Practices for Single-Cell Analysis across Modalities. Nat. Rev. Genet. 2023, 24 (8), 550–572. 10.1038/s41576-023-00586-w.

(8) Domínguez Conde, C.; Xu, C.; Jarvis, L. B.; Rainbow, D. B.; Wells, S. B.; Gomes, T.; Howlett, S. K.; Suchanek, O.; Polanski, K.; King, H. W.; Mamanova, L.; Huang, N.; Szabo, P. A.; Richardson, L.; Bolt, L.; Fasouli, E. S.; Mahbubani, K. T.; Prete, M.; Tuck, L.; Richoz, N.; Tuong, Z. K.; Campos, L.; Mousa, H. S.; Needham, E. J.; Pritchard, S.; Li, T.; Elmentaite, R.; Park, J.; Rahmani, E.; Chen, D.; Menon, D. K.; Bayraktar, O. A.; James, L. K.; Meyer, K. B.; Yosef, N.; Clatworthy, M. R.; Sims, P. A.; Farber, D. L.; Saeb-Parsy, K.; Jones, J. L.; Teichmann, S. A. Cross-Tissue Immune Cell Analysis Reveals Tissue-Specific Features in Humans. Science 2022, 376 (6594), eabl5197. 10.1126/science.abl5197.

(9) Alquicira-Hernandez, J.; Sathe, A.; Ji, H. P.; Nguyen, Q.; Powell, J. E. scPred: Accurate Supervised Method for Cell-Type Classification from Single-Cell RNA-Seq Data. Genome Biol. 2019, 20 (1), 264. 10.1186/s13059-019-1862-5.

(10) Song, D.; Chen, S.; Lee, C.; Li, K.; Ge, X.; Li, J. J. Synthetic Control Removes Spurious Discoveries from Double Dipping in Single-Cell and Spatial Transcriptomics Data Analyses. BioRxiv Prepr. Serv. Biol. 2024, 2023.07.21.550107. 10.1101/2023.07.21.550107.

(11) Zhang, J. M.; Kamath, G. M.; Tse, D. N. Valid Post-Clustering Differential Analysis for Single-Cell RNA-Seq. Cell Syst. 2019, 9 (4), 383–392.e6. 10.1016/j.cels.2019.07.012.

(12) Song, D.; Li, K.; Ge, X.; Li, J. J. ClusterDE: A Post-Clustering Differential Expression (DE) Method Robust to False-Positive Inflation Caused by Double Dipping. Res. Sq. 2023, rs.3.rs-3211191. 10.21203/rs.3.rs-3211191/v1.

(13) Neufeld, A.; Gao, L. L.; Popp, J.; Battle, A.; Witten, D. Inference after Latent Variable Estimation for Single-Cell RNA Sequencing Data. Biostatistics 2023, 25 (1), 270–287. 10.1093/biostatistics/kxac047.

(14) Ball, T. M.; Squeglia, L. M.; Tapert, S. F.; Paulus, M. P. Double Dipping in Machine Learning: Problems and Solutions. Biol. Psychiatry Cogn. Neurosci. Neuroimaging 2020, 5 (3), 261–263. 10.1016/j.bpsc.2019.09.003.

(15) Desaire, H. How (Not) to Generate a Highly Predictive Biomarker Panel Using Machine Learning. J. Proteome Res. 2022, 21 (9), 2071–2074. 10.1021/acs.jproteome.2c00117.

(16) Webb-Robertson, B.-J. M.; Wiberg, H. K.; Matzke, M. M.; Brown, J. N.; Wang, J.; McDermott, J. E.; Smith, R. D.; Rodland, K. D.; Metz, T. O.; Pounds, J. G.; Waters, K. M. Review, Evaluation, and Discussion of the Challenges of Missing Value Imputation for Mass Spectrometry-Based Label-Free Global Proteomics. J. Proteome Res. 2015, 14 (5), 1993–2001. 10.1021/pr501138h.

(17) Lazar, C.; Gatto, L.; Ferro, M.; Bruley, C.; Burger, T. Accounting for the Multiple Natures of Missing Values in Label-Free Quantitative Proteomics Data Sets to Compare Imputation Strategies. J. Proteome Res. 2016, 15 (4), 1116–1125. 10.1021/acs.jproteome.5b00981.

(18) Vanderaa, C.; Gatto, L. Revisiting the Thorny Issue of Missing Values in Single-Cell Proteomics. J. Proteome Res. 2023, 22 (9), 2775–2784. 10.1021/acs.jproteome.3c00227.

(19) Li, M.; Mallikarjun, V.; Frey, A.; Ogundimu, E.; Trost, M. Empirical-Bayes and Bayesian Hierarchical Modelling for Missingness and Differential Expression in Proteomics. Bioinformatics January 15, 2026. 10.64898/2026.01.15.699650.

(20) Goh, W. W. B.; Wang, W.; Wong, L. Why Batch Effects Matter in Omics Data, and How to Avoid Them. Trends Biotechnol. 2017, 35 (6), 498–507. 10.1016/j.tibtech.2017.02.012.

(21) Geman, D.; d’Avignon, C.; Naiman, D. Q.; Winslow, R. L. Classifying Gene Expression Profiles from Pairwise mRNA Comparisons. Stat. Appl. Genet. Mol. Biol. 2004, 3, Article19. 10.2202/1544-6115.1071.

(22) Tan, A. C.; Naiman, D. Q.; Xu, L.; Winslow, R. L.; Geman, D. Simple Decision Rules for Classifying Human Cancers from Gene Expression Profiles. Bioinforma. Oxf. Engl. 2005, 21 (20), 3896–3904. 10.1093/bioinformatics/bti631.

(23) Afsari, B.; Fertig, E. J.; Geman, D.; Marchionni, L. switchBox: An R Package for k-Top Scoring Pairs Classifier Development. Bioinformatics 2015, 31 (2), 273–274. 10.1093/bioinformatics/btu622.

(24) Marzouka, N.-A.-D.; Eriksson, P. multiclassPairs: An R Package to Train Multiclass Pair-Based Classifier. Bioinformatics 2021, 37 (18), 3043–3044. 10.1093/bioinformatics/btab088.

(25) Leduc, A.; Huffman, R. G.; Cantlon, J.; Khan, S.; Slavov, N. Exploring Functional Protein Covariation across Single Cells Using nPOP. Genome Biol. 2022, 23 (1), 261. 10.1186/s13059-022-02817-5.

(26) Montalvo Landivar, A. P.; Gao, Z.; Liu, M.; Gruskin, Z. L.; Leduc, A.; Preza, S.; Xie, Y.; Rozo, A. V.; Ahn, J. H.; Straubhaar, J. R.; Doliba, N.; Stoffers, D. A.; Slavov, N.; Alvarez-Dominguez, J. R. An Adult Clock Regulator Links Circadian Rhythms to Pancreatic β-Cell Maturation. BioRxiv Prepr. Serv. Biol. 2025, 2023.08.11.552890. 10.1101/2023.08.11.552890.

(27) Ai, L.; Binek, A.; Zhemkov, V.; Cho, J. H.; Haghani, A.; Kreimer, S.; Israely, E.; Arzt, M.; Chazarin, B.; Sundararaman, N.; Meyer, J. G.; Sharma, A.; Marbán, E.; Svendsen, C. N.; Van Eyk, J. E. Single-Cell Proteomics Reveals Specific Cellular Subtypes in Cardiomyocytes Derived From Human iPSCs and Adult Hearts. Mol. Cell. Proteomics MCP 2025, 24 (9), 100910. 10.1016/j.mcpro.2025.100910.

(28) Furtwängler, B.; Üresin, N.; Richter, S.; Schuster, M. B.; Barmpouri, D.; Holze, H.; Wenzel, A.; Grønbæk, K.; Theilgaard-Mönch, K.; Theis, F. J.; Schoof, E. M.; Porse, B. T. Mapping Early Human Blood Cell Differentiation Using Single-Cell Proteomics and Transcriptomics. Science 2025, 390 (6770), eadr8785. 10.1126/science.adr8785.

(29) Khan, S.; Elcheikhali, M.; Leduc, A.; Huffman, R. G.; Derks, J.; Franks, A.; Slavov, N. Inferring Post-Transcriptional Regulation within and across Cell Types in Human Testis. BioRxiv Prepr. Serv. Biol. 2024, 2024.10.08.617313. 10.1101/2024.10.08.617313.

(30) Petrosius, V.; Aragon-Fernandez, P.; Arrey, T. N.; Woessmann, J.; Üresin, N.; de Boer, B.; Su, J.; Furtwängler, B.; Stewart, H.; Denisov, E.; Petzoldt, J.; Peterson, A. C.; Hock, C.; Damoc, E.; Makarov, A.; Zabrouskov, V.; Porse, B. T.; Schoof, E. M. Quantitative Label-Free Single-Cell Proteomics on the Orbitrap Astral MS. Mol. Cell. Proteomics MCP 2025, 24 (6), 100982. 10.1016/j.mcpro.2025.100982.

(31) Saddic, L.; Kaneda, G.; Momenzadeh, A.; Zilberberg, L.; Song, Y.; Mastali, M.; Kreimer, S.; Hutton, A.; Haghani, A.; Meyer, J.; Parker, S. Single Cell Proteomics Reveals Novel Cell Phenotypes in Marfan Mouse Aneurysm. BioRxiv Prepr. Serv. Biol. 2025, 2025.02.15.638465. 10.1101/2025.02.15.638465.

(32) Amidan, B. G.; Orton, D. J.; Lamarche, B. L.; Monroe, M. E.; Moore, R. J.; Venzin, A. M.; Smith, R. D.; Sego, L. H.; Tardiff, M. F.; Payne, S. H. Signatures for Mass Spectrometry Data Quality. J. Proteome Res. 2014, 13 (4), 2215–2222. 10.1021/pr401143e.

(33) Pedregosa, F.; Varoquaux, G.; Gramfort, A.; Michel, V.; Thirion, B.; Grisel, O.; Blondel, M.; Prettenhofer, P.; Weiss, R.; Dubourg, V.; Vanderplas, J.; Passos, A.; Cournapeau, D.; Brucher, M.; Perrot, M.; Duchesnay, É. Scikit-Learn: Machine Learning in Python. J. Mach. Learn. Res. 2011, 12, 2825–2830. 10.5555/1953048.2078195.

(34) Rocchi, P.; Ferreri, A. M.; Simone, G.; Bagnara, G. P.; Paolucci, G. Neuronal Cell Differentiation of Human Neuroblastoma Cells by Inducing Agents in Combination. Anticancer Res. 1991, 11 (5), 1885–1889.

